# SUMOylation contributes to proteostasis of the chloroplast protein import receptor TOC159 during early development

**DOI:** 10.1101/2020.07.12.198945

**Authors:** Sonia Accossato, Felix Kessler, Venkatasalam Shanmugabalaji

**Affiliations:** Laboratory of Plant Physiology, University of Neuchâtel, 2000 Neuchâtel, Switzerland

## Abstract

Chloroplast biogenesis describes the transition of non-photosynthetic proplastids to photosynthetically active chloroplasts in the cells of germinating seeds. Chloroplast biogenesis requires the import of thousands of nuclear-encoded preproteins and depends on the essential import receptor TOC159, mutation of which results in non-photosynthetic albino plants. We previously showed that ubiquitin-proteasome system (UPS)-dependent regulation of TOC159 levels contributes to the regulation of chloroplast biogenesis during early plant development. Here, we demonstrate that the SUMO (Small Ubiquitin-related Modifier) pathway crosstalks with the ubiquitin-proteasome pathway to affect TOC159 stability during early plant development. We identified a SUMO3-interacting motif (SIM) in the TOC159 GTPase (G-) domain and a SUMO3 covalent SUMOylation site in the membrane (M-) domain. A single K to R substitution (K1370R) in the M-domain disables SUMOylation. Expression of the TOC159K1370R mutant in the *toc159* mutant (*ppi2*) complemented the albino phenotype. Compared to wild type TOC159, TOC159K1370R was destabilized under UPS-inducing stress conditions. However, TOC159K1370R recovered to same protein level as wild type TOC159 in the presence of a proteasome inhibitor. Thus, SUMOylation partially stabilizes TOC159 against UPS-dependent degradation under stress conditions. Our data contribute to the evolving model of tightly controlled proteostasis of the TOC159 import receptor during proplastid to chloroplast transition.

## Introduction

Chloroplasts are unique organelles that carry out photosynthesis. Although chloroplasts contain their own genome, the majority of chloroplast proteins are encoded by the nuclear genome and synthesized as preproteins in cytosol, and these preproteins are imported into the chloroplast through TOC-TIC complexes (**T**ranslocon at the **O**uter or **I**nner membrane of the **C**hloroplast) [1]. The core of the TOC complex contains two related GTP-dependent preprotein receptor GTPases, Toc159 and Toc33, which interact with a β-barrel membrane protein, Toc75, that forms a protein-conducting channel, and is regulated by specific interactions with nuclear encoded preproteins [2-4]. Toc159 is a major point of entry for highly abundant photosynthesis-associated preproteins arriving at the translocon complex. It is therefore regarded as the major chloroplast protein import receptor. Toc159 has a three-domain structure: a highly acidic N-terminal domain (A-domain), a central GTP-binding domain (G-domain) and a C-terminal membrane anchor domain (M-domain). Chloroplast biogenesis, the transition of a non-photosynthetic proplastid to a photosynthetically active chloroplast, depends on the essential import receptor Toc159 and its mutation results in non-photosynthetic albino plants [5].

Recent studies have shown that during the etioplast to chloroplast transition, the TOC components are ubiquitinated by a novel chloroplast RING-type E3 ubiquitin ligase SP1 (SUPPRESSOR OF PPI1 LOCUS 1). Ubiquitinated TOC components are extracted from the chloroplast outer envelope membrane with the help of SP2 (an Omp85-like β-barrel protein) and Cdc48 (a cytosolic AAA+ chaperone) providing the extraction force. The ubiquitinated TOC-component is degraded by the 26S proteasome in the cytosol [6, 7]. This proteolytic pathway has been named CHLORAD. In a different context, chloroplast biogenesis takes place during the proplastid to chloroplast transition. It is dependent on the plant hormone gibberellic acid (GA). Under unfavorable seed germination conditions when gibberellic concentrations are low, a DELLA (RGL2) (a negative regulator of GA signaling) promotes the ubiquitylation and degradation of Toc159 by the 26S proteasome to insertion into the outer membrane of the chloroplast. This mechanism delays the onset of chloroplast biogenesis at an early developmental stage and has been shown to be independent of SP1 and presumably CHLORAD [8].

A recent study employed a Yeast-two-Hybrid screen to identify putative Arabidopsis SUMO substrates. The E2 SUMO conjugating enzyme (SCE1) was used as bait and TOC159 was identified as a high probability interaction candidate. TOC159 containing a predicted SAS (SUMO Attachment Site – ΨKXE/D) [9]. The Small Ubiquitin-like Modifier (SUMO), a 11-kDa protein, covalently modifies a large number of proteins. SUMO-dependent regulation is involved in many cellular processes, including gene expression, signal transduction, genome maintenance, protein localization and activity [10]. SUMOylation plays a crucial role in plant development and stress responses [11-13]. This reversible and dynamic SUMOylation starts with the attachment of SUMO to the target protein by a conjugation pathway, mechanistically analogous to the ubiquitylation system [12, 14]. The target protein covalently modified by SUMO, performs specific functions, which may subsequently be reversed by SUMO proteases that hydrolyze the isopeptide bond between SUMO and the target protein [15]. In addition, SUMO can also interact with target proteins through a SUMO interacting motif (SIM). The non-covalent SUMO-SIM interaction may work as a molecular signal for protein-protein interaction and affect stability of proteins [16].

In this study, we address the role of Small Ubiquitin Like Modifier (SUMO) modification of TOC159 in the context of the proplastid to chloroplast transition and investigate both covalent SUMOylation and non-covalent SUMO-interaction. The biochemical and genetic evidence showed that the TOC159 G-domain contains a SUMO-interacting motif (SIM) that interacts with the SUMO3 and that SUMOylation of TOC159 at the M-domain is the key regulatory mechanism to protect it from further depletion under low gibberellic acid conditions. The data provides new insight for TOC159 SUMO-binding and SUMOylation demonstrating that SUMOylation positively influences protein stability with regard to the UPS. Thereby SUMOylation contributes to the proteostatic fine-tuning of TOC159 levels during TOC159-dependent chloroplast biogenesis in early plant development.

## Results

### SUMO3 interacts with the TOC159 G-domain and is SUMOylated at the TOC159 M-domain

An earlier study using an *in vitro* SUMOylation assay showed that out of three SUMO isoforms (SUMO1, SUMO2, and SUMO3), only SUMO3 SUMOylated TOC159 [11]. TOC159 has a N-terminal A- (acidic-), a central G- (GTP-binding domain) and a C-terminal M- (membrane) domain [5]. The A-domain is known to be non-essential for Toc159 function and exquisitely sensitive to protease activity [17, 18]. It was excluded from our DNA constructs and only the TOC159 G- and M-domains (TOC159GM) were used. In addition to a covalent SUMOylation site TOC159 may also have SUMO-interaction motifs (SIM). To investigate the possibility of SUMO interaction with TOC159GM, we used the GPS-SUMO prediction algorithm (http://sumosp.biocuckoo.org/online.php) to search for SUMO Interaction Motifs (SIM) in TOC159GM [19]. Based on the search results there is a predicted SIM (VKVLP) in the G-domain that is conserved in other plant species. (**Figure 1A** and **B**). To analyze the physical interaction between TOC159GM and SUMO isoforms, we performed yeast two hybrid assays using TOC159G and TOC159M separately as baits. TOC159G interacted exclusively with SUMO3 but not SUMO1 and SUMO2. None of the SUMO isoforms interacted with TOC159M in the yeast two hybrid assay. (**Figure 1C**).

**Figure 1.**
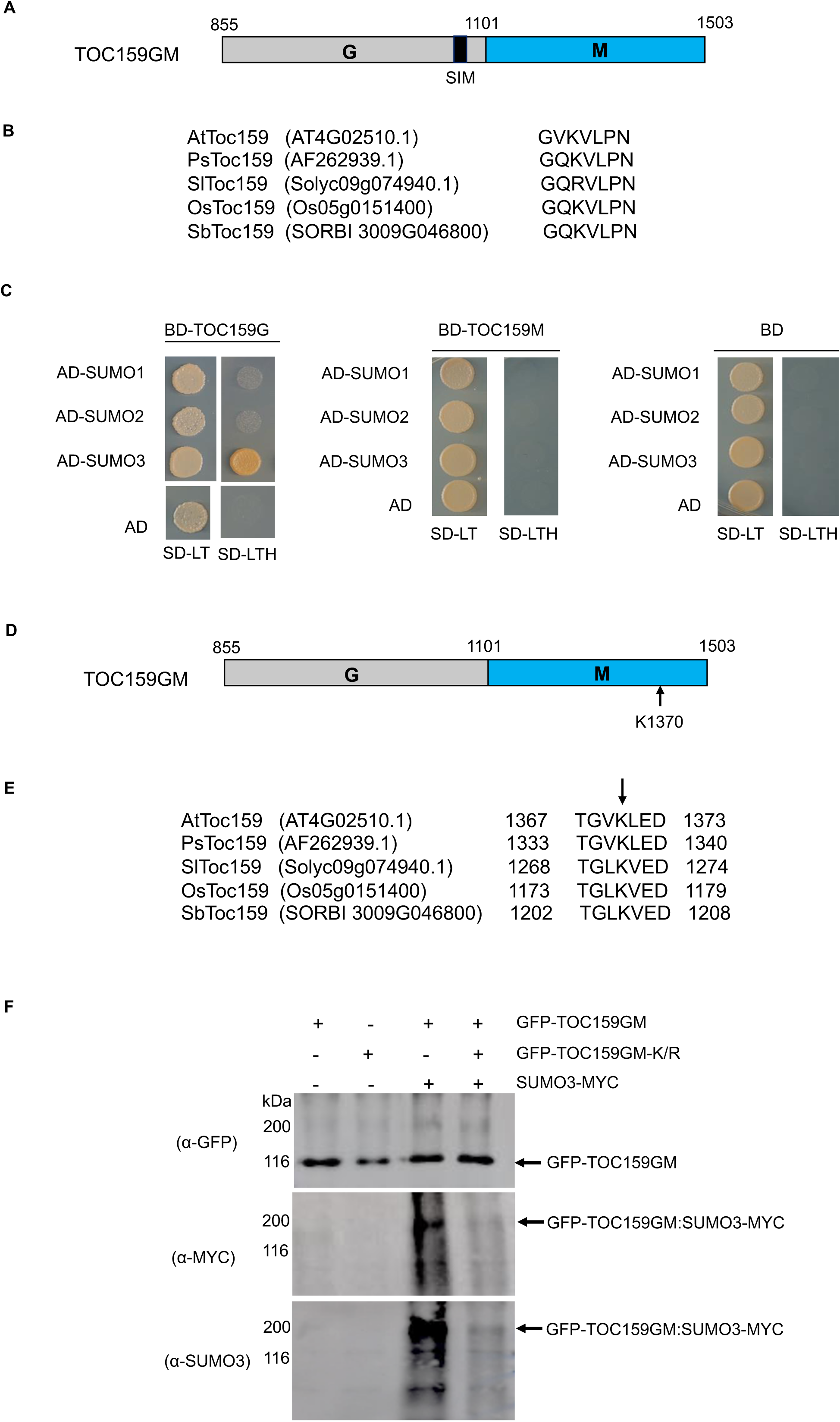
SUMO interaction and SUMOylation of TOC159GM. **(A)** Schematic representation of TOC159GM indicating the predicted SUMO Interaction Motif (SIM) in the G-domain. **(B)** Alignment of putative conserved SIM motifs in the G-domain of the following species: *Arabidopsis thaliana* (At), *Pisum sativum* (Ps), *Solanum lycopersicum* (Sl), *Oryza sativa* (Os), and *Sorghum bicolor* (Sb). **(C)** Yeast two-hybrid interaction assay of TOC159 G and M domain with SUMO proteins on –Leu, –Trp and –Leu, –Trp, –His medium. AD, activation domain; BD, binding domain; empty vector was used as a control. **(D)** Schematic representation of TOC159GM with indication of the predicted SUMOylation site K1370 (Lysine) at the M domain. **(E)** Alignment of the conserved predicted K1370 SUMOylation sites in the M-domain of a variety of species (as in Figure 1B). **(F)** Transient expression of GFP-TOC159GM, GFP-TOC159GM-K/R (SUMO mutant, K1370 replaced with R) with and without SUMO3-MYC in *Nicotiana benthamiana* leaves. Total protein extracts were subjected to immunoprecipitation with anti-GFP beads. The immunoprecipitated proteins from the expression of GFP-TOC159GM (lane 1) and GFP-TOC159GM-K/R (lane 2) alone as well as the co-expression with SUMO3 (lane 3 and 4) were analyzed by western blotting using anti-GFP, anti-MYC and anti-SUMO3 antibodies.

We used the GPS-SUMO algorithm (http://sumosp.biocuckoo.org/online.php) to search for covalent SUMOylation sites in TOC159GM. A high scoring consensus SUMOylation site with a strongly conserved motif (TGVKLED) and containing a potentially SUMOylatable lysine (K1370) was identified within the M-domain. (**Figure 1-Supplement 1**) (**Figure 1D**). The SUMOylation motif as well as K1370 of Arabidopsis are well conserved in other plants species (**Figure 1E**). To investigate the SUMOylation of TOC159GM, we selected the SUMO3 isoform based on the earlier *in vitro* study [9]. We infiltrated *Nicotiana benthamiana* with 35S-GFP-TOC159GM or GFP-TOC159GM-K/R (replacing lysine with a non-sumoylatable arginine residue at position 1370) each together with or without 35S-SUMO3-MYC. To analyze the infiltration experiments identical amounts of total extracts were subjected to immunoprecipitation using anti-GFP-beads followed by Western blotting. An anti-GFP antibody was used to indicate total expression of GFP-TOC159GM and GFP-TOC159GM-K/R and resulted in bands of similar intensities in all four experiments. Anti-MYC and anti-SUMO3 were used to analyze conjugation of SUMO3-MYC to GFP-TOC159GM and GFP-TOC159GM-K/R. The Western blotting using anti-MYC and anti-SUMO3 antibodies resulted in strong signals for GFP-TOC159GM (**Figure 1F**, lane 3) but only very weak ones for GFP-TOC159GM-K/R (**Figure 1F**, lane 4) when co-expressed with SUMO3-MYC. No signals were detected when the GFP-TOC159GM constructs were expressed in the absence of SUMO3-MYC (**Figure** 1F, lanes 1 and 2). Note that GFP-TOC159GM co-infiltration with SUMO3 resulted in a much higher molecular mass band when analyzed with anti-MYC or -SUMO3 than the main, strong GFP-TOC159GM band detected with anti-GFP. It therefore appears that upon co-expression with SUMO3-MYC only a small fraction of GFP-TOC159GM was present in the SUMOylated form and that it had a higher molecular mass than GFP-TOC159GM alone (**Figure 1F**).

### The non-SUMOylatable TOC159GM-K/R mutant complements the *ppi2* mutation

It has been demonstrated that the A-domain of TOC159 is dispensable but that the M-domain of TOC159 is essential for protein import into the chloroplast [20]. Furthermore, it was demonstrated that TOC159GM alone without the A-domain could complement the albino *ppi2* phenotype [17]. To characterize the effect of the K1370R mutation on the M-domain *in vivo*, we engineered transgenic lines expressing GFP-TOC159GM as well as GFP-TOC159GM-K/R under the TOC159 promoter in the *ppi2* background. Two independent transgenic lines of pTOC159-GFP-TOC159GM:*ppi2* and pTOC159-GFP-TOC159GM-K/R:*ppi2* (called GFP-TOC159GM:*ppi2 and* GFP-TOC159GM-K/R:*ppi2* plants hereafter) were isolated and the genotypes were confirmed by PCR using specific primer pairs (**Figure 2A, Figure 2-figure Supplement 1A**). Homozygous GFP-TOC159GM:*ppi2* and GFP-TOC159GM-K/R:*ppi2* transgenic lines gave green plants with almost identical chlorophyll concentrations that were very close to those of the wild type (**Figure 2C**). This indicated complementation of the albino *ppi2* phenotype. Western blotting analysis using GFP antibodies specifically detected GFP-TOC159GM and GFP-TOC159GM-K/R proteins. The blots revealed that TOC159GM and GFP-TOC159GM-K/R proteins accumulated to very similar levels in the respective transgenic lines. The blots were also probed with antibodies against TOC75 and TOC33, showing no significant differences between the two transgenic lines (**Figure 2B**) (**Figure 2-figure supplement 1B and C**). To localize the GFP-TOC159GM and GFP-TOC159GM-K/R *in vivo* 7-day-old seedlings were analyzed by confocal fluorescence microscopy Both GFP-TOC159GM and GFP-TOC159GM-K/R gave clear fluorescent signals at the chloroplast periphery that are consistent with outer envelope membrane localization (**Figure 2D**).

**Figure 2.**
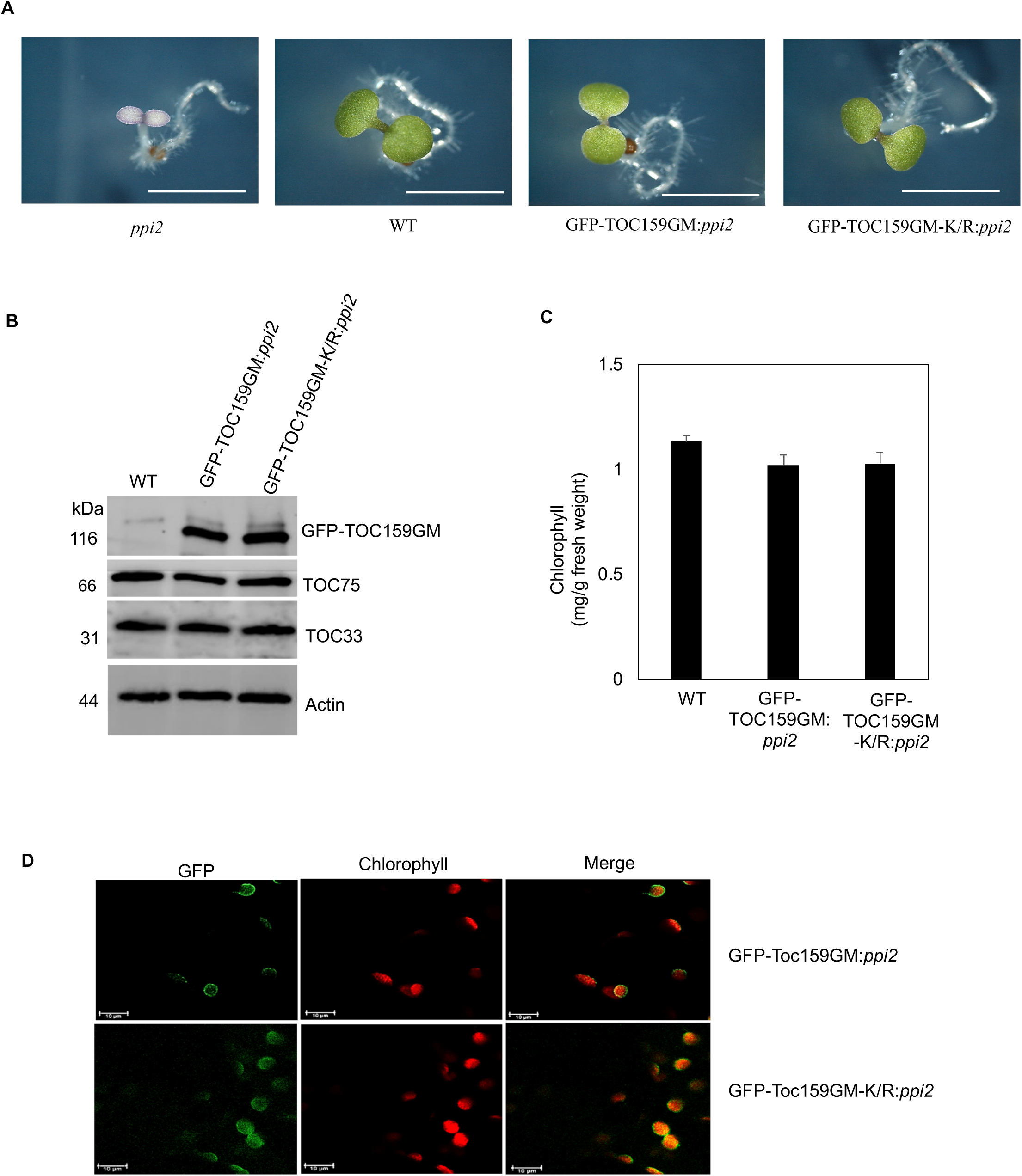
Complementation of *ppi2* (*toc159* mutant) by non-SUMOylatable TOC159GM-K/R. **(A)** Phenotypes of 7 days old seedling of *ppi2*, WT (Ws), GFP-TOC159GM:*ppi2* and GFP-TOC159GM-K/R:*ppi2*. **(B)** Western blot analysis of total protein extracts from 7 days old seedlings of WT (Ws), GFP-TOC159GM:*ppi2* and GFP-TOC159GM-K/R:*ppi2*. The blot was probed with anti-GFP, -TOC75 and TOC33 antibodies. Actin was used as a loading control. **(C)** Chlorophyll levels of wild type, GFP-TOC159GM:*ppi2* and GFP-TOC159GM-K/R:*ppi2* from 7 days old seedlings. **(D)** Confocal microscopy analysis of 7 days old GFP-TOC159GM:*ppi2* and GFP-TOC159GM-K/R:*ppi2* seedlings. Green fluorescence (GFP, left-hand panel), red fluorescence (Chlorophyll, middle pane) and the overlay of the two (right-hand panel) are shown.

### TOC159GM-K/R accumulation is diminished when compared to TOC159GM under low gibberellic acid conditions

Addition of paclobutrazol (PAC) to the MS growth medium can be used to inhibit GA biosynthesis and results in low GA conditions [21]. Under low GA during early germination, the DELLA (RGL2) destabilizes TOC159 via the ubiquitin-proteasome system [8]. To explore a possible role of SUMOylation of K1370 on protein stability GFP-TOC159GM:*ppi2* and GFP-TOC159GM-K/R:*ppi2* were allowed to germinate in the presence or absence of PAC. As reported earlier, the GFP-TOC159GM protein level was severely reduced in GFP-TOC159GM:*ppi2* seeds under low GA. In comparison, the level of GFP-TOC159-K/R in GFP-TOC159GM-K/R:*ppi2* seeds was even more diminished. Moreover, TOC75 and TOC33 levels were also lower in mutant TOC159GM-K/R:*ppi2* than in wildtype GFP-TOC159GM:*ppi2* under low GA whereas their levels were the same in the untreated lines (**Figure 3A and B**) (**Figure 3 - figure supplement 1 and 2**). PAC-treated seeds, low in GA, accumulated very high levels of the RGL2 protein [21]. We also compared RGL2 protein levels between TOC159GM:*ppi2* and TOC159GM-K/R:*ppi2* in the presence of PAC. The results revealed that there was no difference in RGL2 accumulation between the two lines (**Figure 3C**).

**Figure 3.**
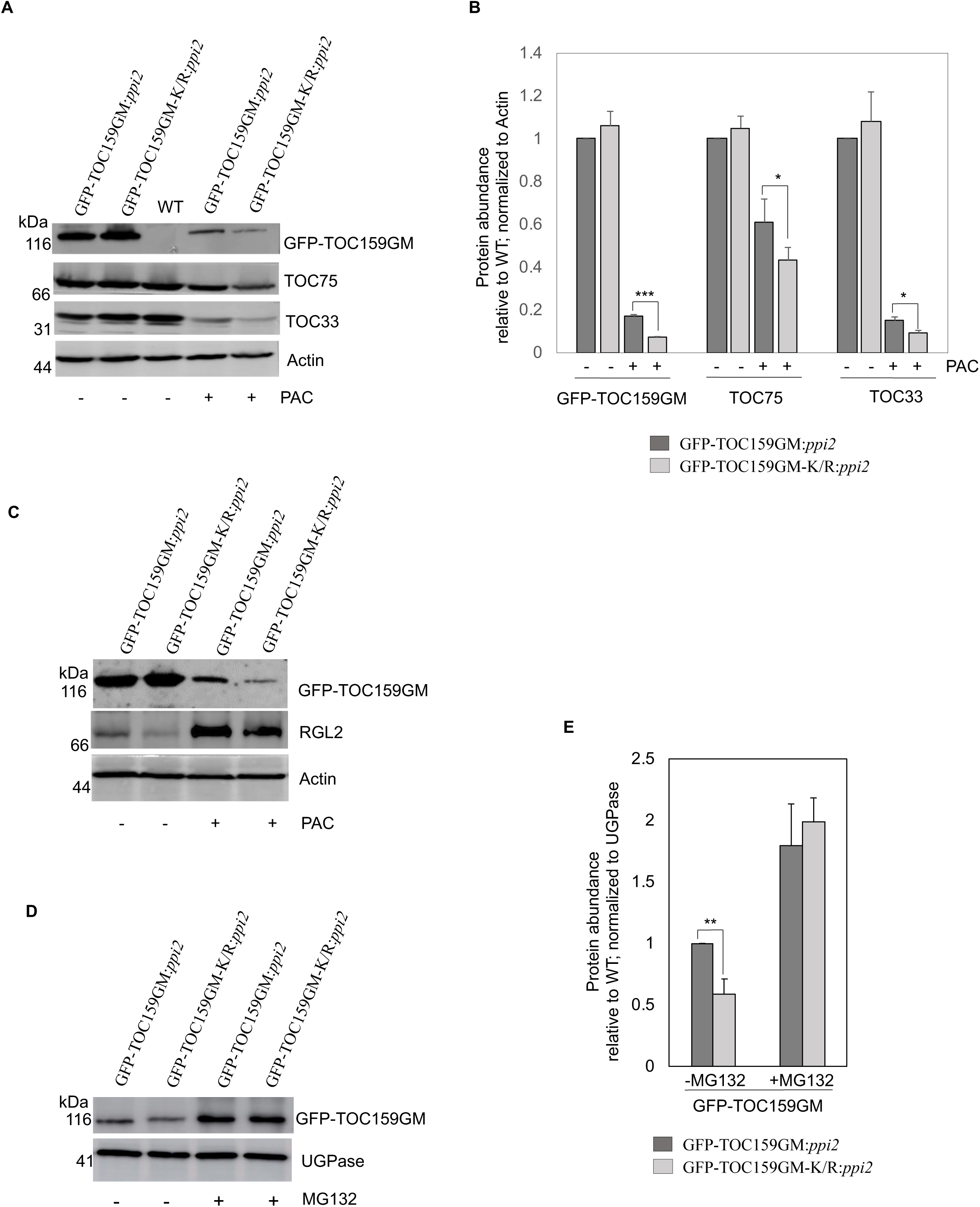
SUMOylation partially protects TOC159 from UPS-mediated degradation. **(A)** Immunoblotting of total protein extracts from three days old seedling of GFP-TOC159GM:*ppi2* and GFP-TOC159GM-K/R:*ppi2* grown in the presence or absence of PAC (5 µM). WT (Ws) was used as the control for antibody specificity. The blot was probed with anti-GFP, -TOC75, and -TOC33 antibodies. Anti-actin was used for a loading control. **(B)** Specific bands corresponding to GFP, TOC75, TOC33 and actin were quantified (A). The quantified bands were normalized to GFP-TOC159GM in GFP-TOC159GM:*ppi2* plants grown in the absence of PAC. Error bars indicate ± SEM (n = 3). Student’s t test; *p < 0.05; ***p < 0.005. **(C)** Immunoblotting of total protein extracts from three days old GFP-TOC159GM:*ppi2* and GFP-TOC159GM-K/R:*ppi2* seedlings grown in the presence or absence of PAC (5 µM). The blot was probed with anti-GFP and -RGL2 antibodies. Anti-actin was used for a loading control. **(D)** Total protein extracts of three days old GFP-TOC159GM:*ppi2* or GFP-TOC159GM-K/R:*ppi2* grown seedling grown on PAC and subsequently treated with or without MG132 were analyzed by immunoblotting using anti-GFP antibodies and anti-UGPase for a loading control. **(E)** The specific bands corresponding to GFP and UGPase were quantified (C). The quantified bands were normalized to GFP-TOC159GM in GFP-TOC159GM:*ppi2* without MG132. Error bars indicate ± SEM (n = 3). Student’s t test; **p < 0.01.

### SUMOylation partially stabilizes the TOC159 under low gibberellic acid conditions

To determine whether the diminished stability of GFP-TOC159GM-K/R under low GA in the presence of PAC is also due to UPS-mediated degradation, GFP-TOC159GM-K/R:*ppi2* and GFP-TOC159GM:*ppi2* seeds were germinated in the presence of PAC and subjected to treatment with or without the proteasome inhibitor MG132. Western blot analysis demonstrated that both GFP-TOC159GM-K/R and GFP-TOC159GM were rescued by MG132 and accumulated to the same level. (**Figure 3E and F**) (**Figure 3 - figure supplement 3**). The results suggest that SUMOylation partially protects GFP-TOC159GM against UPS-mediated degradation under low GA when compared to GFP-TOC159GM-K/R.

### Accumulation of photosynthesis-associated proteins trended lower in TOC159GM-K/R:ppi2 than TOC159GM:ppi2

The results so far demonstrated that non-SUMOylatable TOC159GM-K/R is significantly more susceptible to UPS-mediated degradation than wildtype TOC159GM under low GA. To address whether this has an effect on the accumulation of photosynthesis-associated proteins (the presumed TOC159 cargo proteins), western blotting was carried out comparing GFP-TOC159GM-K/R:*ppi2* to GFP-TOC159GM:*ppi2.* It has been shown that the expression of most photosynthesis-associated genes did not change significantly in the presence of moderate and low PAC concentrations (5 and 1 μM) [22]. To eliminate the possibility of gene expression effects on protein accumulation, GFP-TOC159GM:*ppi2* and GFP-TOC159GM-K/R:*ppi2* seedlings were grown in the presence or absence of an intermediate 2μM PAC concentration. The western blot analysis revealed that, the accumulation of the two photosynthesis-associated proteins (PSBO1 and PSBA) systematically trended lower but not to statistically significant extent in TOC159GM-K/R:*ppi2* when compared to TOC159GM:*ppi2* (**Figure 3 - Figure Supplement 4**).

## Discussion

It has been demonstrated previously that TOC159 physically interacts with the SUMO E2 enzyme in a yeast two-hybrid screen and that the SUMO3 isoform covalently SUMOylated TOC159 in an *in vitro* assay [9]. Apart from a covalent SUMOylation motif, the GPS-SUMO algorithm identified a conserved non-covalent SUMO-interacting motif (SIM) (“VKVLP”) in the G-domain of TOC159. We confirmed this prediction using a yeast two hybrid assay. It also revealed that the G-domain specifically interacted with the SUMO3 isoform and not with SUMO1 and -2. (**Figure 1D**). It has been shown in Arabidopsis that the GA receptor GID1 interacted with SUMO1 through a SIM that prevented the itsinteraction with a DELLA protein [23]. It is tempting to hypothesize that non-covalent binding of TOC159(SIM) to SUMO3 may modifyinteractions with SUMOylated complex partners such as TOC33/TOC75 or the DELLA RGL2 under low GA conditions as these proteins also have predicted SUMOylation sites (http://sumosp.biocuckoo.org/online.php).

Toc159 has highly conserved SUMOylation motif (“TGV**K**LED”) with a lysine at position 1370 (K1370) within the M-domain of TOC159. We established SUMOylation at K1370 using an *in planta* SUMOylation assay in *N. benthamiana* (**Figure 1F**). Expression of non-SUMOylatable GFP-TOC159GM-K/R (containing the K1370R substitution) restored a wildtype green phenotype in the GFP-TOC159GM-(K/R):*ppi2* Arabidopsis plants. Note that *ppi2* mutant plants normally have an albino phenotype. Based on the presence of wildtype levels of TOC75 and TOC33 it appeared that the TOC complex assembled normally in the GFP-TOC159GM-(K/R):*ppi2* plants (**Figure 2A and B**). Confocal laser microscopy localized GFP-TOC159GM-(K/R) at the envelope of the chloroplast (**Figure 2D**). Taken together, these results indicate that outer membrane insertion, Toc complex assembly as well as chloroplast biogenesis take place normally under standard growth conditions despite the K1370R mutation and disabled SUMOylation.

In Arabidopsis, SUMOylation is triggered by environmental stimuli including biotic and abiotic stress. Hormone signaling (GA, auxin, brassinosteroids, ABA) and development (root, seed, embryo and meristem) are the main biological processes associated with SUMOylation in plants. [23-29]. While SUMOylation is independent from the ubiquitination pathway, there is the complex crosstalk between these two pathways. SUMOylation can promote the UPS-dependent degradation through SUMO-targeted Ub ligases (STUBLs) [30]. Contrary to this, SUMOylation may also protect from UPS-dependent degradation by blocking the ubiquitination of lysine residues [31-34]. Here, we explored the connection between TOC159 SUMOylation and the UPS-mediated TOC159 degradation under low GA during early developmental stages in seed germination.

We previously demonstrated that low GA concentrations brought about by paclobutrazol promote the DELLA (RGL2)-dependent TOC159 degradation via the UPS [8]. Under low GA, GFP-TOC159GM-K/R accumulated to significantly (< 50%) lower levels than wildtype GFP-TOC159GM in the respective overexpression lines. In addition, the TOC159-interacting TOC-complex core proteins TOC75 and -33 also accumulated to considerably lower levels in the GFP-TOC159GM-K/R:*ppi2* line under low GA presumably to maintain complex stoichiometry (**Figure 3A and B)**. Typically, a small fraction of the SUMO target protein pool undergoes SUMOylation under a specific cellular condition [35]. The reduced levels under low GA of GFP-TOC159GM-K/R when compared to the wildtype protein were attributed to increased UPS-mediated protein degradation as both proteins recovered to the same protein levels in the presence of the proteasome inhibitor MG132 (**Figure 3D and E**).

We conclude that SUMOylation partially stabilizes TOC159 against UPS-dependent degradation under specific conditions such as low GA during early development. The photosynthesis-associated proteins are considered the preferred transport cargoes of TOC159 because they fail to accumulate in the *ppi2* mutant lacking TOC159. They tended to accumulate to a lesser degree in GFP-TOC159GM-K/R:*ppi2* plants under low GA in the presence of PAC, but not in statistically significant fashion. The explanation may be that the levels of TOC159 are already low in the GFP-TOC159GM-K/R:ppi2 plants in the absence of PAC and allow only for relatively small changes in the accumulation of photosynthesis-associated proteins in the presence of PAC. Nevertheless, the results suggest that SUMOylation at the M-domain of TOC159 serves to fine tune preprotein import under low GA.

Based on the results we propose a hypothetical model for the role of SUMOylation during early developmental stages, when environmental conditions are unfavorable and GA concentrations are low. As previously demonstrated, the chloroplast import receptor TOC159 is ubiquitylated prior to outer membrane insertion by an unknown E3-ligase other than SP1 and degraded via the UPS. In addition, its G-domain interacts with SUMO3, the physiological consequences of which remain unknown but may regulate interaction with DELLA system. The M-domain may be SUMOylated by SUMO3 protecting to some extent against UPS-dependent degradation. When the environmental conditions become more favorable, GA levels increase, and the GA receptor-DELLA complex is degraded by the UPS and TOC159 liberated for outer membrane insertion. The non-ubiquitinated and de-SUMOylated TOC159 is assembled into the TOC complex thus allowing proplastids to differentiate into chloroplasts (**Figure 4**). In the CHLORAD pathway cytosolic Cdc48 extracts ubiquitinated TOC proteins from the outer membrane of the chloroplast [7] [36]. The intriguing connection between the Cdc48 and SUMO pathways in chromatin dynamics [37], suggests that the SUMO pathway might also act on the CHLORAD pathway at specific developmental stage but we currently do not offer evidence for such a mechanism. Our data specifically implicate SUMOylation and probably SUMO-interaction in the control of proplastid to chloroplast transition by regulation of TOC159 levels in early plant development.

**Figure 4.**
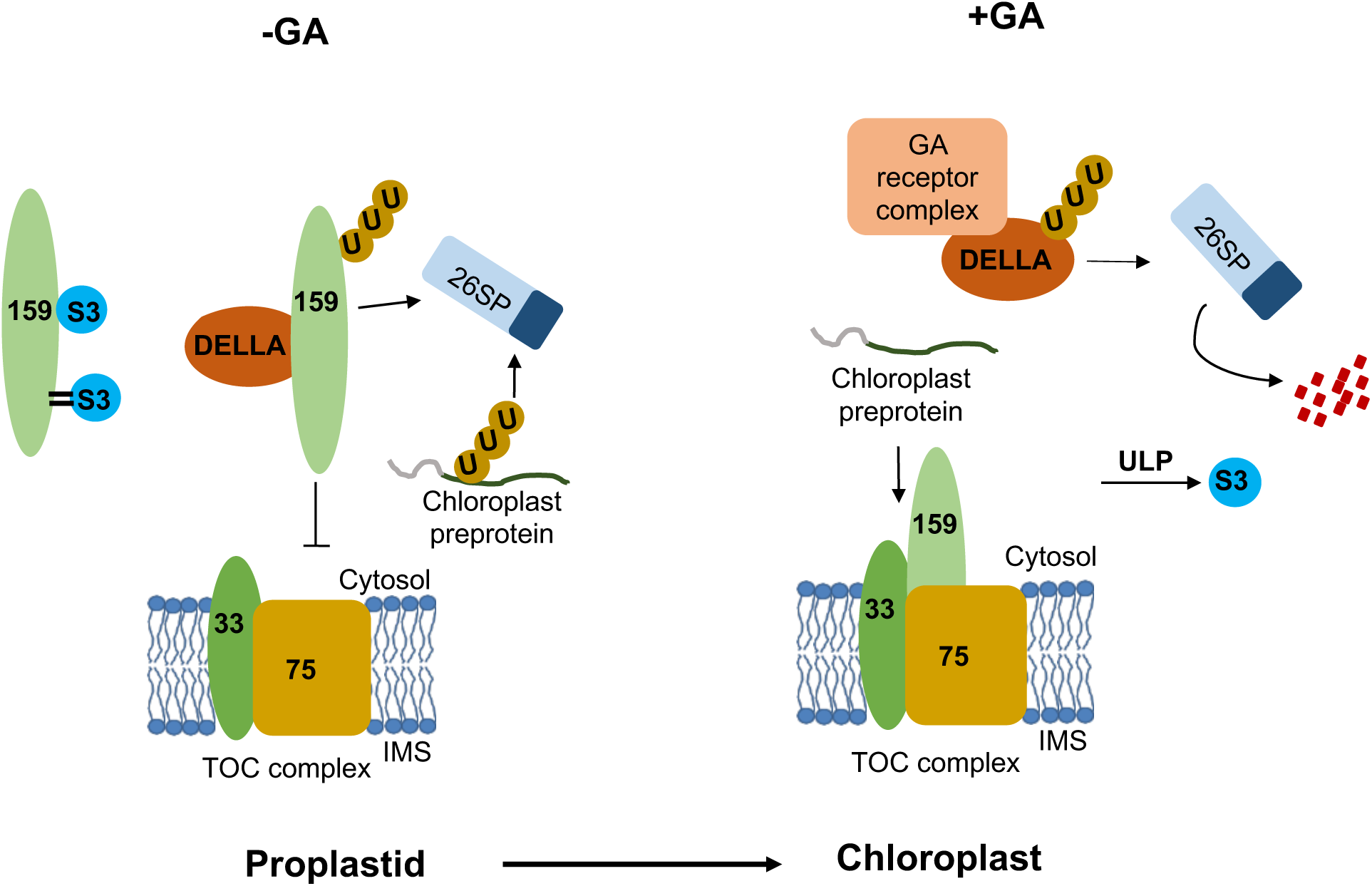
Hypothetical model for the SUMOylation dependent fine-tuning of chloroplast biogenesis at the level of the TOC159 import receptor in early plant development. Environmental conditions influence the concentrations of GA that accumulate upon seed imbibition. When active GA levels are reduced by stress (-GA; left hand panel), DELLA (RGL2) accumulates and sequesters TOC159, which is then degraded via the UPS. In addition, TOC159 interacts and is also covalently SUMOylated by SUMO3. Covalent SUMOylation protects TOC159 against UPS-mediated degradation and supports the accumulation of photosynthesis-associated proteins in the chloroplast. Any un-imported preproteins are degraded in the cytosol via the UPS. When GA concentrations increase (+GA, right hand panel), the GA receptor complex binds to DELLA, which is degraded by the UPS. TOC159 is then free to assemble into the TOC complex. Presumable, deSUMOylation of TOC159 by a SUMO protease (ULP) assists assembly into the TOC complex. Protein import is thus enabled, allowing proplastids to differentiate into chloroplasts

## Acknowledgements

This work was supported by grants from the Swiss National Science Foundation (31003A_156998 and 31003A _176191) and by the University of Neuchâtel.

## Figure Legends

**Figure 1 Supplement 1.**
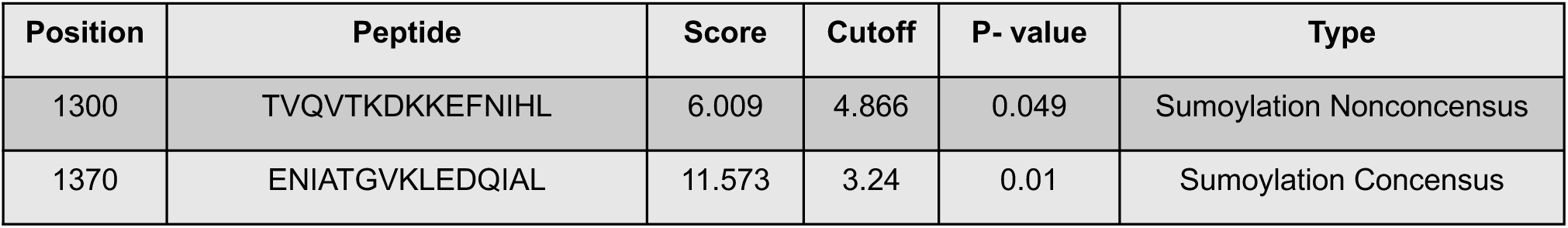
Predicted SUMOylation sites at TOC159GM. Predicted SUMOylation sites at TOC159GM domain using the GPS SUMO prediction algorithm with a high threshold (http://sumosp.biocuckoo.org/online.php).

**Figure 2 Supplement 1.**
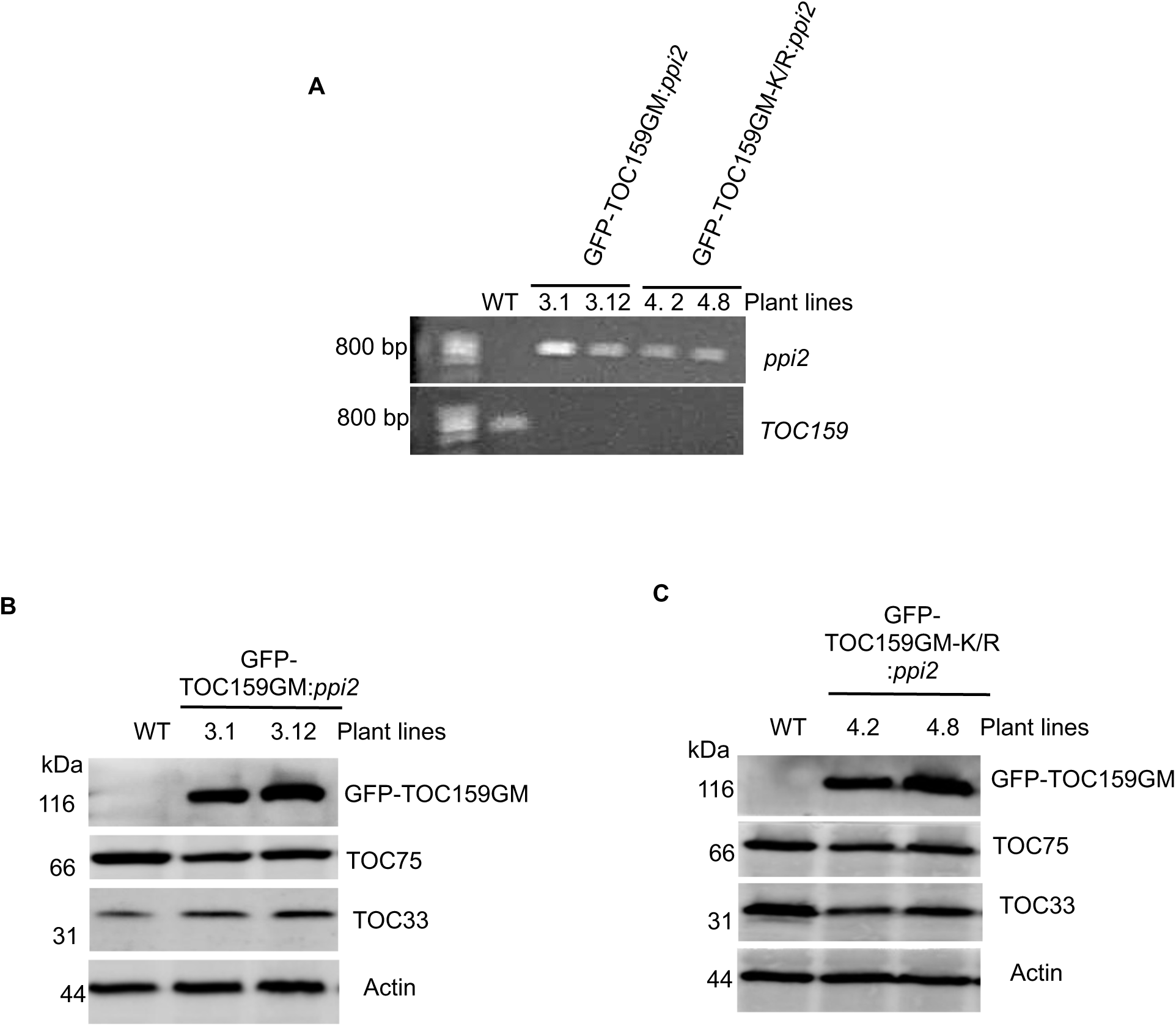
Screening of *ppi2* plants for complementation with GFP-TOC159GM and GFP-TOC159GM-K/R. **(A)** Genotyping of two independent plant lines (3.1 and 3.12) expressing GFP-TOC159GM in the *ppi2* background (GFP-TOC159GM:*ppi2*) and two independent plant lines (4.2 and 4.8) expressing GFP-TOC159GM-K/R in the *ppi2* background (GFP-TOC159GM-K/R:*ppi2*). WT was used as the control for primers. The *ppi2* and TOC159 primer sets were used. **(B)** Immunoblotting of total protein extracts of seedlings from two independent plant lines (3.1 and 3.12) expressing GFP-TOC159GM in the *ppi2* background (GFP-TOC159GM:*ppi2*). WT was used as the control for antibody specificity. The following antibodies were used GFP, TOC75, and TOC33 proteins. Actin was used as a loading control. **(C)** Immunoblotting of total protein extracts of seedlings from two independent plant lines (4.2 and 4.8) expressing GFP-TOC159GM-K/R in *ppi2* background (GFP-TOC159GM-K/R:*ppi2*). WT was used as the control for antibody specificity. The following antibodies were used GFP, TOC75, and TOC33 proteins. Actin was used as a loading control.

**Figure 3 Supplement 1.**
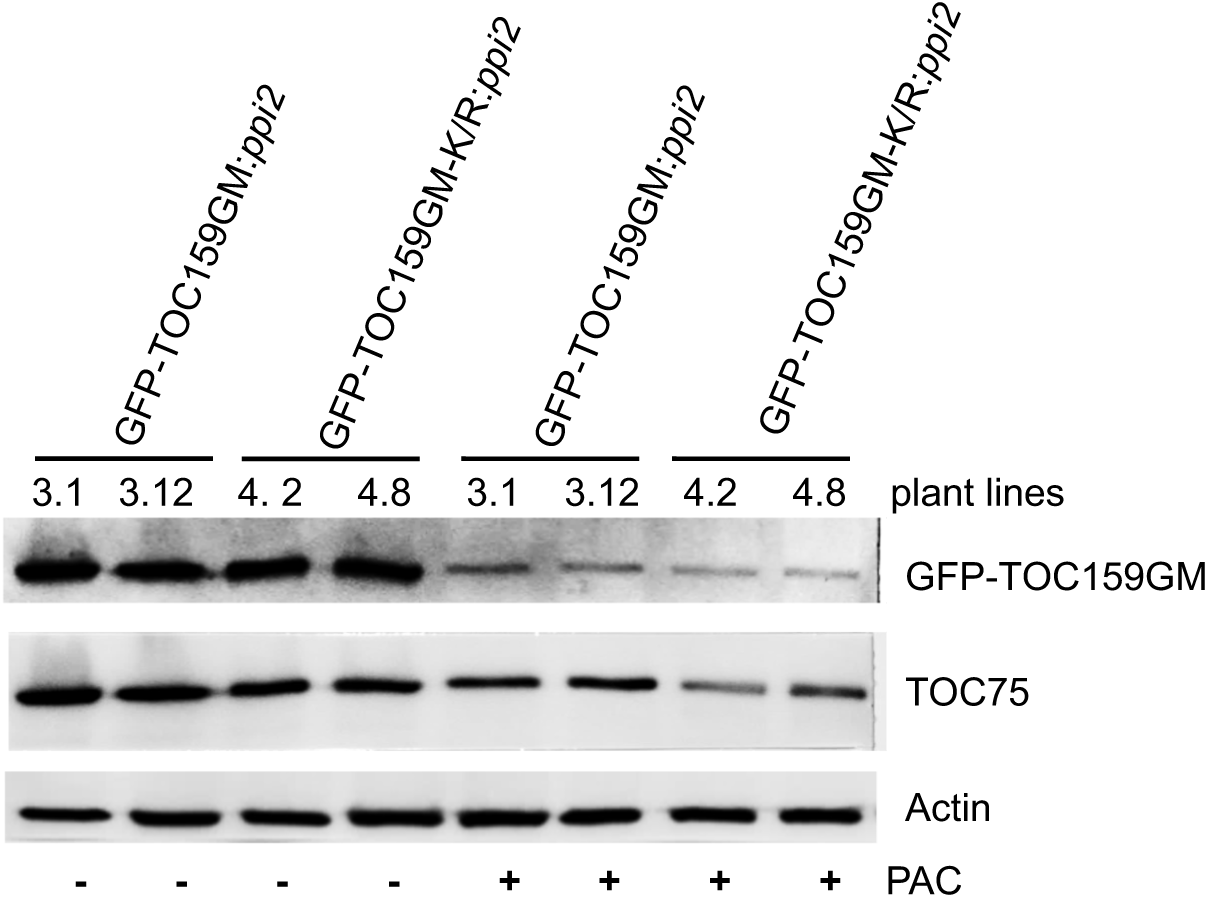
Immunoblotting of total protein extracts from two independent GFP-TOC159GM:*ppi2* lines (3.1 and 3.12) and two independent GFP-TOC159GM-K/R:*ppi2* lines (4.2 and 4.8). Total protein was extracted from three days old seedlings grown in the presence or absence of PAC (5 µM). WT was used as the control for antibody specificity. The following antibodies were used GFP, TOC75, and TOC33 proteins. Actin was used as a loading control.

**Figure 3 Supplement 2.**
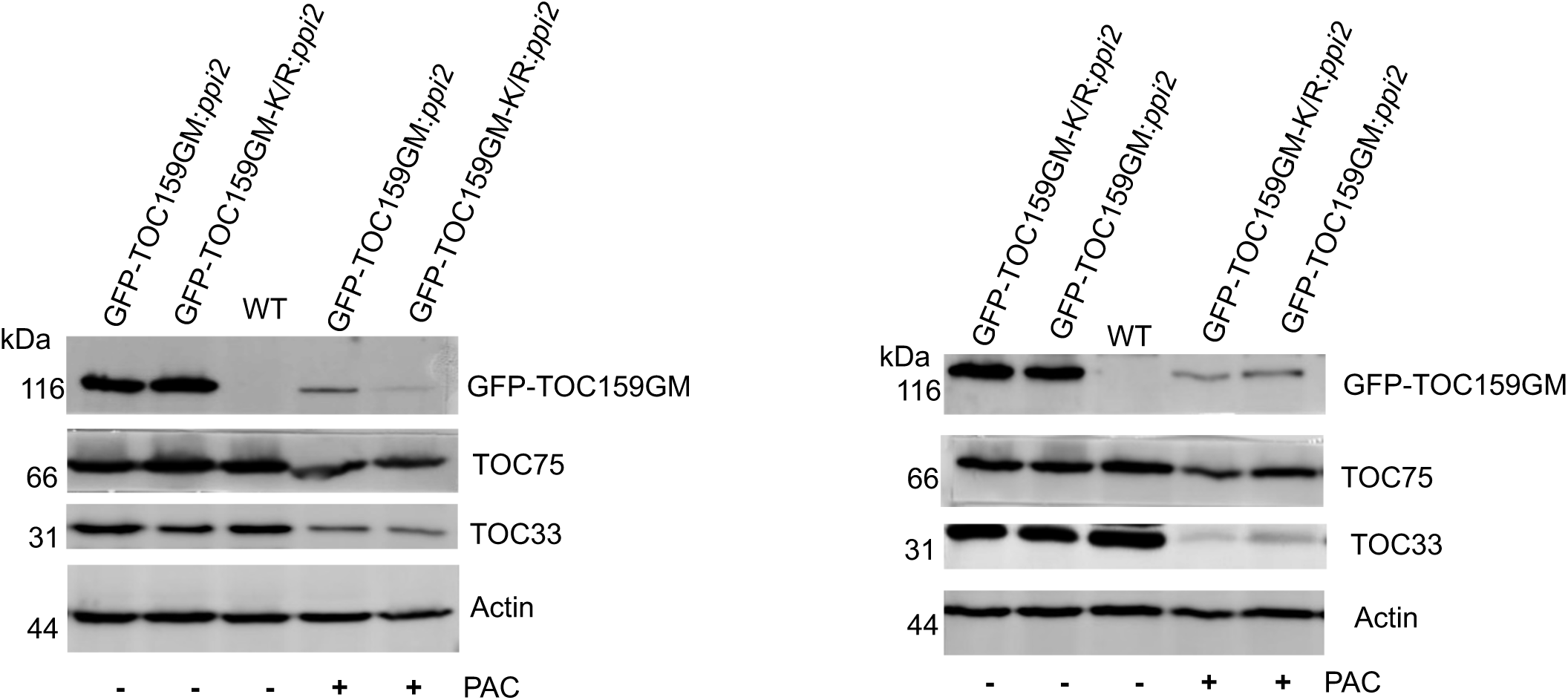
Immunoblotting of total protein extracts from GFP-TOC159GM:*ppi2* and GFP-TOC159GM-K/R:*ppi2* grown in the presence or absence of PAC (5 µM). WT was used as the control for antibody specificity. The following antibodies were used GFP, TOC75, and TOC33 proteins. Actin was used as a loading control. The specific bands corresponding to GFP, TOC75, TOC33 and actin were quantified. The quantified bands were normalized to GFP-TOC159GM in GFP-TOC159GM:*ppi2* grown in the absence of PAC. These data were used in figure 3B.

**Figure 3 Supplement 3.**
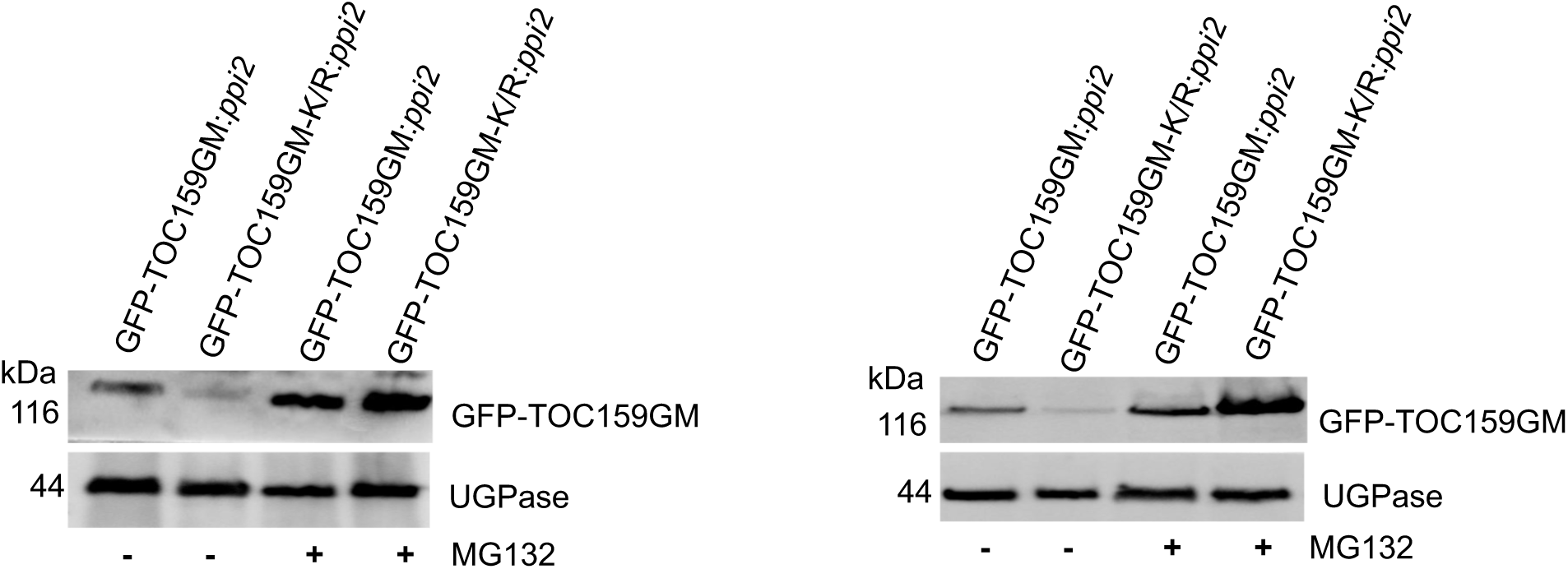
Total protein extracts of three days old GFP-TOC159GM:*ppi2* or GFP-TOC159GM-K/R:*ppi2* grown in the presence of PAC and further treated with or without MG132, analyzed by immunoblotting using antibodies against GFP. UGPase was used as a loading control. The specific bands corresponding to GFP and UGPase were quantified. The quantified bands were normalized to GFP-TOC159GM in GFP-TOC159GM:*ppi2* without MG132. These data were used for figure 3E.

**Figure 3 Supplement 4.**
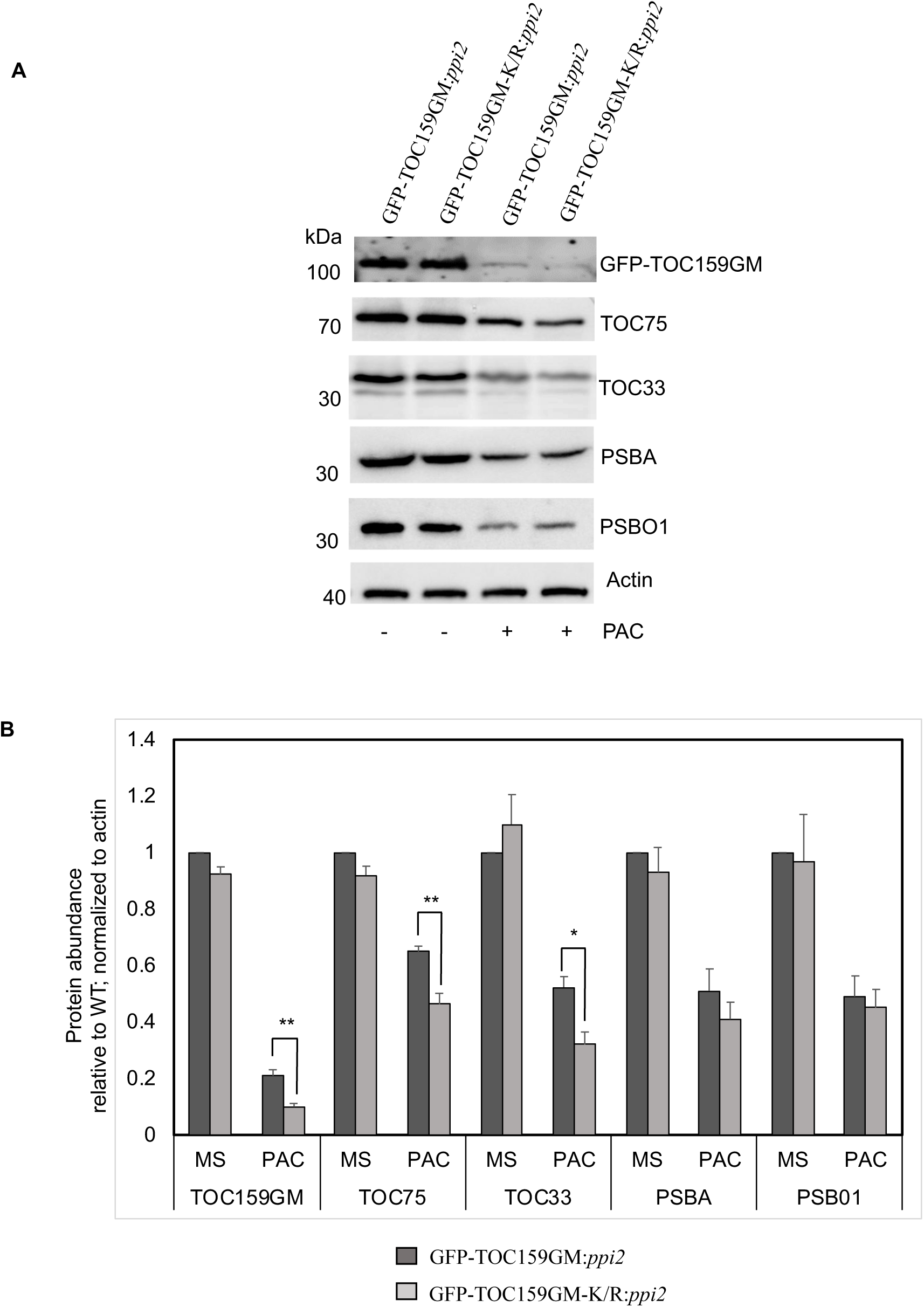
The TOC159 SUMOylation-deficient line TOC159GM-K/R:*ppi2* accumulates two photosynthesis-associated proteins trended lower compared to TOC159GM:*ppi2* under low GA conditions. **(A)** Immunoblotting of total protein extracts from three days old seedling of GFP-TOC159GM:*ppi2* and GFP-TOC159GM-K/R:*ppi2* grown in the presence or absence of PAC (2 µM). The blot was probed with anti-GFP, -TOC75, - TOC33, -PSBA and -PSBO1 antibodies. Anti-actin was used as a loading control. **(B)** The specific bands corresponding to GFP, TOC75, TOC33, PSBA, PSBO1 and actin were quantified (A). The quantified bands were normalized to GFP-TOC159GM in GFP-TOC159GM:*ppi2* grown in the absence of PAC. Error bars indicate ± SEM (n = 3). Student’s t test; *p < 0.05; **p < 0.01.

## List of supplementary tables

**Supplementary file1:** List of primers used in this study

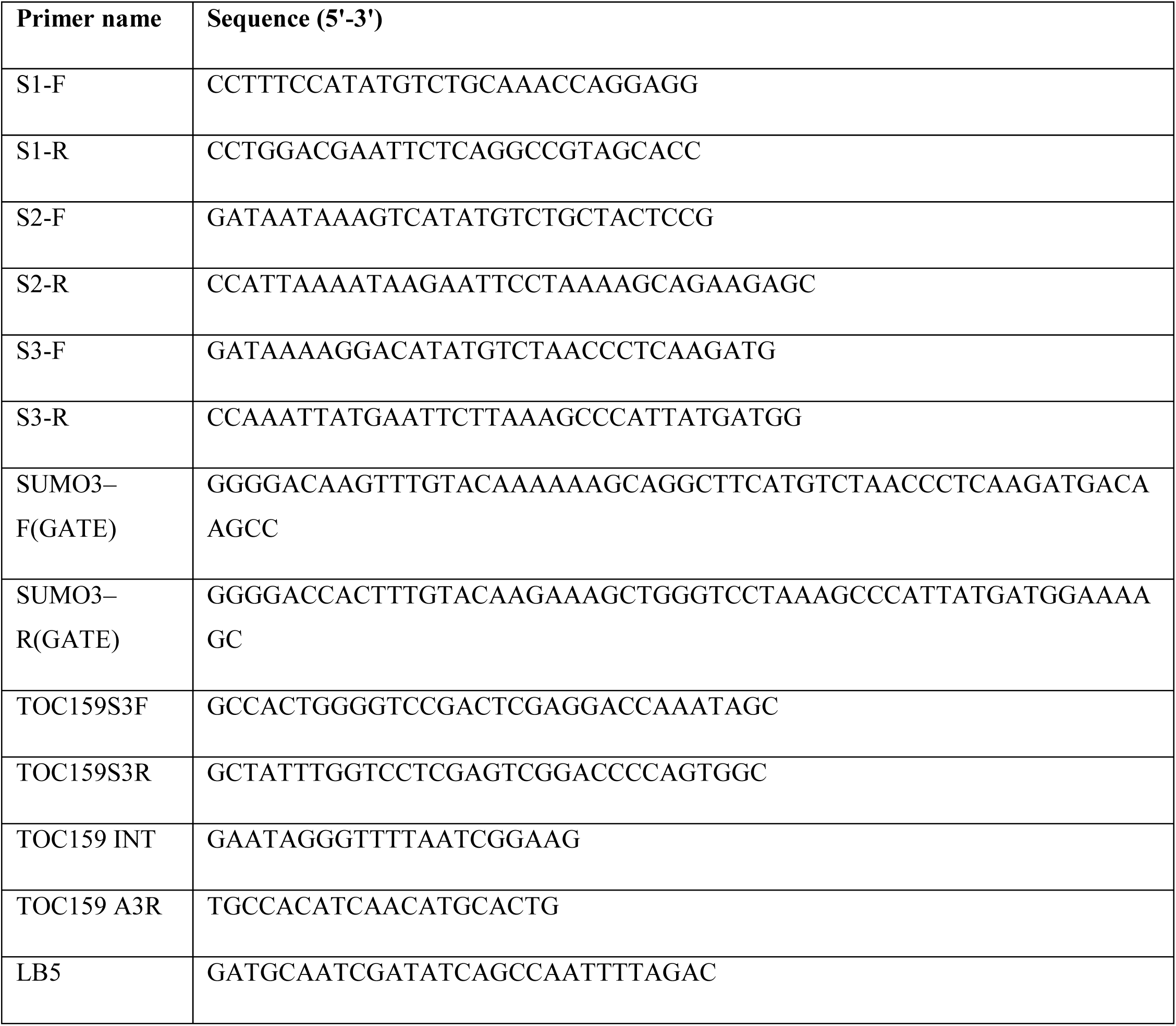

## Materials & Methods

### Plant materials and growth conditions

The *A. thaliana* Columbia- Wassilewskija (Ws) ecotype was used as wild types. The *ppi2* mutant and pTOC159-GFP-TOC159GM:*ppi2* line used in this study were in the Wassilewskija (Ws) ecotype previously described [5])[8]. Seedlings were grown on Murashige-Skoog (MS) medium with long-day conditions (16-h light, 8-h dark, 120 µmol x m-2 x s-1, 21 °C). Plants were grown either on soil under short-day conditions (8 hours dark, 16 hours light, 120 µmol x m-2 x s-1, 21 °C) for vegetative growth, and under long-day conditions (16-h light, 8-h dark, 120 µmol x m-2 x s-1, 21 °C) for flower development and seed production.

### Seedling treatment

Surface-sterilized seeds were placed on MS medium supplemented with 2 μM or 5 μM paclobutrazol (PAC). The plates were incubated under short-day conditions for 3 days. Proteasome inhibitor experiments were performed as described earlier [8].

### Plant transformation and transgenic lines

The K1370R point mutation was introduced into the binary construct pTOC159-GFP-TOC159GM using a site directed mutagenesis kit (Agilent-QuikChangeII) with the primers TOC159S3F and TOC159S3R and resulted in pTOC159-GFP-TOC159GM-K/R. The pTOC159-GFP-TOC159GM-K/R construct was introduced into *Agrobacterium tumefaciens* (C58 strain) and stably transformed into heterozygous *ppi2* plants using the floral dip method. Transformed plants were selected on phosphinothricin-containing medium and lines homozygous for the transgene as well as the *ppi2* mutation were isolated and named pTOC159:GFP-TOC159GM-K/R:*ppi2* (referred to as GFP-TOC159GM-K/R:*ppi2* plants).

### Yeast two-hybrid assays

The pGBKT7-TOC159G (BD fusion) and pGBKT7-TOC159M (BD fusion) vector were introduced in to the yeast strain Y2H GOLD as previously described [8]. The full-length cDNA sequences of SUMO1, SUMO2, and SUMO3 were amplified using primers (S1F, S1R, S2F, S2R, S3F, S3R), digested with NdeI/ EcoRI and ligated into the corresponding sites of the pGADT7 vector. The empty bait vector (BD) was used as a negative control. Co-transformants were selected on SD –Leu –Trp and SD –Leu –Trp –His plates.

### Confocal laser scanning microscopy

One week old seedlings of the GFP-TOC159GM:*ppi2* and GFP-TOC159GM-K/R:*ppi2* lines were directly observed under a Leica TCS SL confocal microscope. Fluorescence images were captured and analyzed using LCS lite software (Leica).

### Protein extraction and immunoblotting

Identical amounts of samples were collected and proteins extracted using AP extraction buffer (100 mM Tris pH 8, 2% b-mercaptoethanol, 4% SDS, 20% glycerol) followed by acetone precipitation [38]. The SDS-PAGE and immunoblotting were performed according to standard protocols. To probe the blots, primary antibodies recognizing TOC159 [5], TOC75 [39], TOC33 [40], GFP (Takara), SUMO3 (Agrisera), UGPase (Agrisera), actin (sigma) and RGL2 [21] were used. As markers for photosynthesis-associated proteins, antibodies recognizing PSBA and PSBO1 were purchased from Agrisera. Secondary antibodies were anti-rabbit IgG conjugated with horseradish peroxidase (Millipore), or goat anti-mouse IgG conjugated with horseradish peroxidase (Sigma). Chemiluminescence was detected using ECL Plus Western Blotting Detection Reagents (Pierce) and developed using a GE Amersham Imager 600. Band intensities were quantified using ImageQuant TL (GE Healthcare) software.

### *In planta* CoIP from transient expression system in *N. benthamiana*

Full-length SUMO3 was PCR amplified from cDNA using the primers SUMO3–F(GATE) and SUMO3–R2(GATE) and inserted into the pENTR221 vector by BP clonase (Invitrogen). It was recombined into the pEarleyGate 203 binary vector with LR clonase to obtain a 35S-SUMO3-MYC fusion construct. The binary vector 35S-GFP-TOC159GM used in this study has been described previously (Shanmugabalaji *et al.*, 2018). The point mutation was introduced in the binary construct 35S-GFP-TOC159GM by using a site-directed mutagenesis kit (Agilent-QuikChangeII) with the primers TOC159S3F and TOC159S3R, as results we obtained 35S-GFP-TOC159GM-K/R. The 35S-SUMO3-MYC, 35S-GFP-TOC159GM, and 35S-GFP-TOC159GM-K/R were introduced into *Agrobacterium tumefaciens* (C58 strain). And co-infiltrated into 23 weeks old *N. benthamiana*. Immunoprecipitation to isolate the protein complexes from total protein extracts using GFP-tagged microbeads (Miltenyi Biotec) has been described previously [8]. Anti-GFP antibody were used to detect GFP-TOC159, GFP-TOC159GM-(K/R). Anti-SUMO3, and anti-MYC antibodies, respectively, were used to identify the SUMOylated TOC159GM.

### Quantification and statistical analysis

For protein quantification on immunoblots, ImageQuant TL (GE Healthcare) software was used to measure band intensities, and the data shown as mean ± SEM. Statistical analysis was carried using the Student’s t-test, with p values higher than 0.05 being considered non-significant (n.s.) while p values lower than 0.05 being considered significant for the analyzed data and indicated as: *, p < 0.05; **, p < 0.01; ***, p < 0.005.

